# Can Individual Internal Models Predict Idiosyncratic Scene Exploration?

**DOI:** 10.64898/2026.04.01.715777

**Authors:** Micha Engeser, Nasibeh Babaei, Daniel Kaiser

**Affiliations:** Neural Computation Group, Department of Mathematics and Computer Science, Physics, Geography, Justus Liebig University Giessen, 35392 Giessen, Germany; Center for Mind, Brain and Behavior (CMBB), Philipps University Marburg, Justus Liebig University Giessen, and Technical University Darmstadt, 35032 Marburg, Germany; Center for Applied Computer Science and Data Science (ZAD), Justus Liebig University Giessen, 35392 Giessen, Germany; Cluster of Excellence “The Adaptive Mind”, Justus Liebig University Giessen, Philipps University Marburg, and Technical University Darmstadt, 35392 Giessen, Germany

## Abstract

Each individual person looks at natural scenes in their own unique way, resulting in a distinct perceptual experience of the world. However, little is known about why such differences in gaze emerge. Here, we test the hypothesis that idiosyncrasies in gaze behavior are predicted by inter-subject variations in internal models—expectations about how scenes typically look. In two experiments, we first characterized participants’ personal internal models by asking them to draw typical bathroom and kitchen scenes. Individual differences in these drawings were quantified using an objective deep learning pipeline and, in turn, related to individual differences in gaze behavior. In Experiment 1, where participants freely viewed a set of kitchen and bathroom photographs, inter-subject similarities in internal models did not predict inter-subject similarities in gaze. In Experiment 2, we encouraged strategic exploration through gaze-contingent viewing and a memory task. Here, inter-subject similarities in internal models predicted similarities in fixation frequency and the sequence in which different object categories were inspected. These findings suggest that the influence of internal models on visual exploration is stronger under increased sensory uncertainty and when expectation-guided sampling of the environment is encouraged. Together, our results provide new insights into how individual expectations shape gaze behavior and help explain why people differ in how they explore the visual world.

## Introduction

When two individuals view the same scene, one may assume that their perceptual experience is largely identical. However, a growing body of research suggests otherwise: Individuals systematically differ in how they describe the same scene (Coco & Keller, 2012; Kollenda et al., 2025; Yun et al., 2013), how scenes are represented in the brain (Borovska & de Haas, 2024; Charest et al., 2014; Engeser & Kaiser, 2025; Han & Bonner, 2026; Lee & Geng, 2017) and how scene information guides their behavior (Engeser & Kaiser, 2025; Richler et al., 2019; Vanmarcke & Wagemans, 2016; Wang, Chen, et al., 2025; Wang et al., 2024).

Particularly striking individual differences emerge in eye movements during natural scene exploration (Andrews & Coppola, 1999; Bargary et al., 2017; Broda & de Haas, 2024; de Haas et al., 2019; Hayes & Henderson, 2017; Henderson & Luke, 2014; Kollenda et al., 2025). For example, individuals reliably differ in their general eye movement dynamics (e.g. how often they inspect new objects or return to previously inspected ones; Linka & de Haas, 2026) or spatial biases (e.g., how often they fixate the upper or lower parts of objects; Broda & de Haas, 2024). Moreover, individuals also differ in the salience of semantic categories, with some observers preferentially fixating faces and others are consistently drawn to text or motion cues (de Haas et al., 2019). Despite this evidence, we still know relatively little about *why* people differ in where they look when viewing the same image.

Some evidence suggests that genetic factors partially shape gaze behavior (Constantino et al., 2017; Kennedy et al., 2017). At the same time, viewing patterns change across development, becoming less idiosyncratic and more structured into adulthood (Linka et al., 2025). This developmental trajectory suggests that gaze behavior is shaped by accumulated experience. Because individuals grow up in different environments and are exposed to distinct “visual diets” (Hartley, 2022; Mollon et al., 2017), differences in visual experience may ultimately contribute to why people explore scenes differently.

Our lifelong experiences with certain types of environments are thought to give rise to internal models for these environments—expectations about how they typically look like (Engeser et al., 2025). Predictive processing accounts of vision posit that these internal models bias perceptual inference and guide information sampling by directing attention toward behaviorally relevant features (Bakst & McGuire, 2021; Friston et al., 2010, 2012; Goettker et al., 2021; Hayhoe et al., 2012; Henderson, 2017; Sclar et al., 2020; Võ et al., 2019). In this framework, gaze behavior is not merely stimulus-driven but reflects an active process of expectation-guided sampling based on internal models of the world.

We therefore hypothesized that individuals with more similar internal models would explore scenes in more similar ways. To test this idea, we used an inter-subject representational similarity analysis (IS-RSA) approach (Finn et al., 2020) to examine whether similarities in internal models predict similarities in gaze behavior. To characterize the content of internal models, we let participants draw typical exemplars of two indoor scene categories, which we used as pictorial descriptors of their internal models (Engeser et al., 2025; Wang, Chen, et al., 2025; Wang et al., 2025; Wang et al., 2024). We reasoned that the similarity of typical scene drawings across participants reflects similarities in the internal models of these participants, allowing us to create a similarity space that reflects how much individuals agree on their internal models for a given scene category (Engeser & Kaiser, 2025). Next, we asked whether inter-subject similarities in internal models are correlated to idiosyncratic gaze behaviors when exploring an independent set of real-world photographs of these scene categories. We quantified individual exploration patterns regarding where participants looked in a scene, how many fixations they made, and the duration and sequence with which different object categories were inspected. We examined the relationship of individual internal models and idiosyncratic scene exploration across two experiments that varied in task demands as well as sensory uncertainty. In Experiment 1, participants freely viewed scene images, while in Experiment 2, they performed a memory encoding task in a gaze-contingent viewing regime, encouraging more strategic exploration.

## Results

We investigated whether individual differences in internal models predict how participants explore natural scenes. To this end, we used an inter-subject representational similarity analysis (IS-RSA) framework, testing whether pairwise similarities in participants’ internal models predict pairwise similarities in their gaze behavior during scene viewing. Both internal models and gaze patterns were therefore quantified as inter-subject representational dissimilarity matrices (IS-RDMs), capturing the similarity structure across all pairs of participants. This allows testing the covariance of participants across different measures (i.e., internal models and gaze behavior).

Internal-model IS-RDMs were characterized through a drawing task (Figure 1a) in which participants drew their most typical versions of bathroom and kitchen scenes. Pairwise similarities between participants’ drawings were quantified using a deep-learning pipeline following Engeser & Kaiser (2025). Specifically, photorealistic versions of the drawings were generated using commercial Draw3D software and presented to a deep neural network (VGG16). Late-layer (fc7) feature representations were extracted and correlated for each pair of participants, yielding similarity estimates between their drawings organized in an IS-RDM capturing individual differences in internal models. To account for variability related to drawing ability or style, we additionally included a control drawing condition in which participants copied identical photographs of scenes, generating a control IS-RDM that was subsequently partialled out in the analyses.

**Figure 1.**
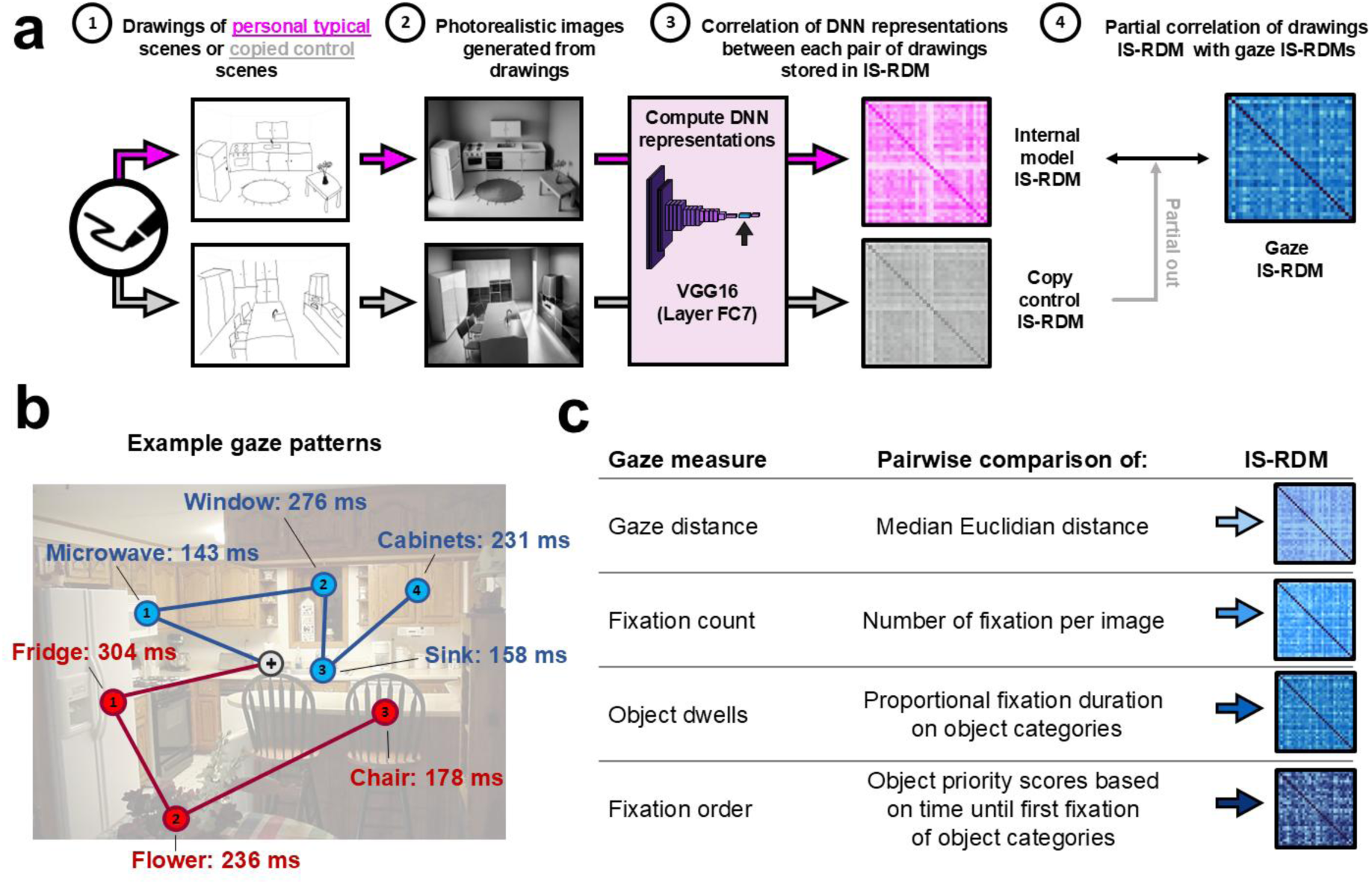
**Inter-subject RSA and gaze measures**. **(a)** A schematic illustration of the inter-subject RSA analysis. First, participants drew a typical version of a scene category (i.e., a kitchen) as a pictorial descriptor of their internal models. Then, drawings were converted to photorealistic images using the Draw3D software and presented to a deep neural network (VGG-16 trained on ImageNet). Correlations of late-layer (fc7) feature activations between each pair of drawings provided similarity estimates of these drawings organized in an IS-RDM. Next, these internal model IS-RDMs were used to predict the similarity structures in gaze IS-RDMs, quantifying individual differences in gaze behavior during exploration of natural scenes. Using partial correlations, we controlled for inter-subject similarities in drawing style, obtained in a copy control condition in which all participants copied the same scene photographs. **(b)** Visualization of the hypothetical gaze patterns of two observers (blue and red). Circles indicate fixation position and lines represent saccades. **(c)** Measures to quantify inter-subject similarities in gaze behavior. Gaze distance: pairwise distance of the median Euclidean distance between observers. Fixation count: pairwise correlations of the number of fixations per image. Object dwells: pairwise correlations of proportional dwell times on selected object categories. Fixation order: pairwise correlations of priority scores (rank-ordered time of first fixation) for a set of selected object categories.

Gaze IS-RDMs were obtained from two eye-tracking experiments with independent groups of participants (n = 34 each) viewing 300 bathroom and kitchen scenes for three seconds. In the free-viewing experiment (Experiment 1), participants explored scenes without specific task demands. In the gaze-contingent experiment (Experiment 2), detailed visual information was restricted to a small window centered on fixation, and participants were instructed to encode scenes for a subsequent memory test, encouraging more strategic exploration.

To quantify inter-subject similarities in gaze behavior, we computed four gaze-based IS-RDMs (Figure 1b-e). First, we computed the median Euclidean distance between participants’ gaze positions. Second, to capture differences in the dynamics of exploration behavior, we further constructed IS-RDMs based on the number of fixations per image. Lastly, two object-based analyses revealed how semantic information is sampled from a scene by measuring the dwell time on different object categories and the order in which these object categories were fixated. Together, these four matrices captured complementary aspects of gaze dynamics and served as outcome measures in the IS-RSA analyses predicting gaze similarity from similarities in internal models.

### Experiment 1: Free viewing

In Experiment 1, participants were asked to freely view the scene images without any specific task.

Before establishing how individual differences in internal models relate to individual differences in gaze, we tested whether individual differences in gaze were reliable in the first place. We used split-half reliability analyses to measure whether inter-subject similarities generalize across different stimulus subsets by correlating IS-RDMs built on two independent halves of the images. Significant positive split-half correlations were observed for all four measures (Figure 2a; gaze distance: *r*=0.95; fixation count: *r*=0.39; object dwells: *r*=0.46; fixation order: *r*=0.40; all *p*<0.001), indicating that we were able to capture reliable individual differences in scene exploration. However, when predicting these individual differences with inter-subject variations in individuals’ internal models, no significant correlations were observed (Figure 2a; gaze distance: r=0.04; fixation count: r<0.01; object dwells: r=-0.01; fixation order: r=-0.03; all p>0.65). Lastly, we assessed whether the order in which objects are drawn in the scene drawings reveals information about the attentional priority of these objects during scene viewing. If so, the correlation of a participant’s drawing order with their own fixation order should be higher than the correlation with other participants’ fixation order. No significant difference between correlations with own and other participants’ fixation order was observed (Δr=-0.01, p=0.71).

**Figure 2.**
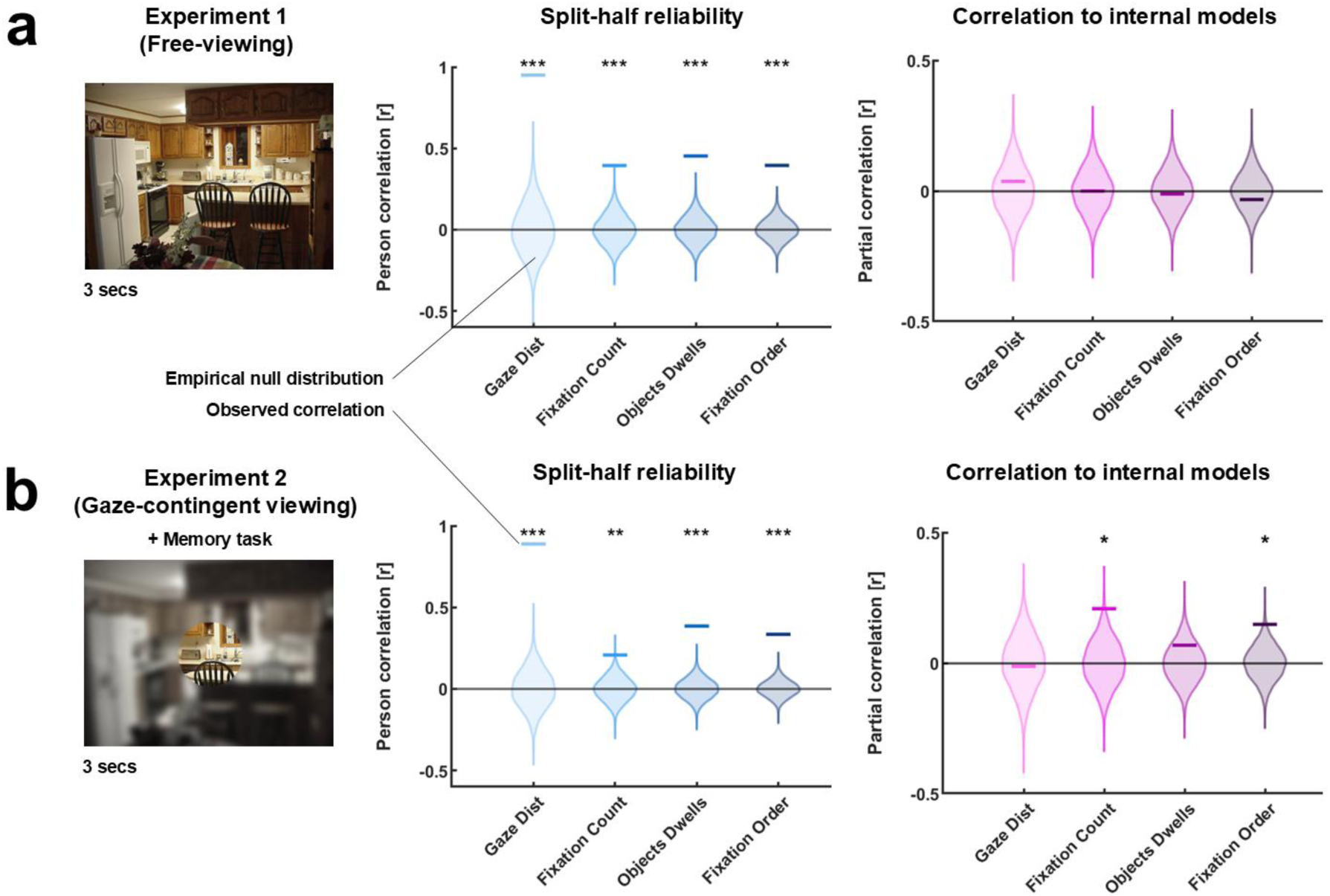
**Inter-subject RSA results**. **(a)** Results from Experiment 1 in which participants could freely view real-world photographs of bathroom and kitchen scenes presented for three seconds. In the split-half reliability analysis, correlations between IS-RDMs built on two independent stimulus subsets revealed reliable individual differences in all gaze measures. Partial correlation of gaze IS-RDMs with IS-RDMs based on participants’ internal models was not significant for any gaze measure. **(b)** Results from Experiment 2, in which detailed scene information was restricted to a small gaze-contingent window to encourage more systematic scene exploration. Additionally, participants were asked to memorize all the scenes for a later memory task. Split-half correlations again demonstrated reliable individual differences for all gaze measures. Inter-subject similarities in the number of fixations per image and fixation order were significantly correlated with inter-subject similarities in internal models. No such relationship was found for the gaze distance and object dwell times. Horizontal bars indicate observed correlation, and violin plots depict the empirical null distribution from a permutation test. * p<0.05, ** p<0.01, and *** p<0.001.

### Experiment 2: Gaze-contingency viewing

In Experiment 2, we encouraged more systematic scene exploration by increasing uncertainty using gaze-contingent viewing, where detailed scene information was restricted to a small window around the current gaze position. Additionally, participants were asked to memorize the scenes for a later memory test. As participants could rely less on peripheral information, they needed to employ their prior knowledge more strongly when moving their gaze around the image, trying to predict where to find relevant information in the image, relative to the current patch’s content.

As in Experiment 1, split-half reliability revealed reliable individual differences (Figure 2b, gaze distance: *r*=0.89, *p*<0.001; fixation count: *r*=0.21, *p*=0.002, object dwells: *r*=0.38, *p*<0.001; fixation order: *r*=0.33, *p*<0.001). When comparing inter-subject similarities across experiments, we found that for all four measures, the median similarity was larger for individuals within the same experiment than for individuals from different experiments (Figure S1; gaze distance: Δ*d*=0.007; fixation count: Δ*r*=0.02; object dwells: Δ*r*=0.01; fixation order: Δ*r*=0.03; all *p*<0.001), indicating that our gaze measures were affected by the gaze contingency condition.

In contrast to Experiment 1, individual differences in internal models correlated with individual differences in gaze (Figure 2b). This was evident in the number of fixations per image (*r*=0.21, *p*=0.025) as well as in the fixation order of selected object categories (*r*=0.15, *p*=0.025). However, gaze distance (*r*=-0.01, *p*=0.55) and object dwell times (*r*=0.07, *p*=0.25) showed no such relationship. When assessing the relationship between drawing and fixation order, we observed that participants’ drawing order was more strongly and positively correlated with their own fixation order than with other participants’ fixation order (Δ*r*=0.03), yet only as a statistical trend (*p*=0.081).

## Discussion

Previous research has demonstrated reliable individual differences in how people explore natural scenes, yet the origins of these differences remain insufficiently understood. Here, we tested the hypothesis that idiosyncrasies in gaze behavior are partly rooted in individual differences in expectations about how scenes are structured. Using inter-subject similarities in drawings as pictorial descriptors of participants’ internal scene models, we show that individual differences in fixation dynamics and object prioritization during scene exploration can partly be predicted from participants’ personal internal models. Crucially, this relationship emerged only when systematic scene exploration was encouraged through gaze-contingent viewing and a memory task but was absent under free-viewing conditions despite reliable individual differences in gaze behavior.

Eye movements are often explained with saliency-based models (Itti et al., 1998; Itti & Koch, 2001), which have been proven successful in predicting the probabilistic distributions of where observers look based on objective image properties. However, they struggle to account for dynamic scan paths and the strategic extraction of meaning from scenes. These aspects are better explained by models incorporating top-down predictive processes alongside bottom-up saliency (Henderson, 2017; Malcolm & Henderson, 2010; Schwetlick et al., 2023). In predictive coding frameworks, eye movements are conceptualized as an active sampling process in which gaze is directed towards locations expected to be maximally informative given prior beliefs and contextual expectations (Bakst & McGuire, 2021; Friston et al., 2010, 2012; Sclar et al., 2020). Our findings align with this view by demonstrating that individual internal models shape how observers sample visual information from complex environments.

Interestingly, the influence of internal models was conditional on the availability of sensory information. When detailed visual information was restricted to a gaze-contingent window, participants were likely encouraged to engage in more strategic exploration, relying on prior expectations to guide information sampling in service of the subsequent memory task. Predictive accounts propose that top-down influences become particularly important when sensory input is limited or ambiguous (de Lange et al., 2018; Henderson, 2017; Malcolm & Henderson, 2010; Peelen et al., 2024; Spotorno et al., 2014). The emerging effect of individual priors under these conditions therefore supports the idea that expectations guide gaze behavior most strongly when bottom-up information is insufficient to momentarily resolve uncertainty.

Notably, internal models predicted fixation frequency and order, but not overall gaze distance or dwell time. Spatial gaze similarity can arise from multiple sources, making this measure less sensitive to expectation-driven influences. For example, gaze distance and object-based fixation analysis may produce diverging results when observers fixate different instances of the same object category or distinct objects in close proximity. The number of fixations a participant performs per image may reflect how well this image aligns with their internal model, whereby higher frequencies of sampling new information serve to reduce uncertainty in less predictable environments (Brunyé & Gardony, 2017; Najemnik & Geisler, 2005, 2008; Rothkegel et al., 2018). Similarly, the order in which objects are fixated provides a window into how internal models prioritize diagnostic scene elements under time constraints. For object dwell times, the slightly positive trend was not significant. Potential explanations could be that object fixation durations were not as strongly impacted by the gaze-contingency manipulation compared to fixation count and order (as indicated by the smaller differences between within and across experiments inter-subject correlations) or that the statistical power was insufficient to detect an effect.

At the same time, reliable individual differences in free-viewing behavior were not predicted by internal models. This is somewhat surprising in the light of recent findings that fixation patterns are best explained by facilitating scene understanding rather than being primarily driven by saliency, also during free-viewing conditions (Murlidaran & Eckstein, 2025). This suggests that gaze idiosyncrasies under unconstrained viewing may be driven by other subject-specific traits, such as biases toward specific image features (Broda & de Haas, 2024; de Haas et al., 2019), visual aesthetic preferences (Ekinci & Kaiser, 2026; Palmer et al., 2013; Vessel et al., 2018), or action affordances (Bartnik et al., 2025; Greene et al., 2016). Alternatively, our drawing-based approximation of internal models may not capture all aspects of internal models that influence gaze behavior: for instance, low-level expectations may be related to color or texture features that were not captured by line drawings (Singer et al., 2023).

Although fixation order was linked to internal models, drawing order was not idiosyncratically aligned with gaze order in a significant manner. This discrepancy may reflect strategic differences in drawing strategies. Our design choice of providing 30 seconds for planning the drawing and liberal drawing time restrictions (4 min.) may encourage drawing strategies unrelated to perceptual priorities, such as constructing scenes from structural anchors rather than from the most informative and diagnostic elements. Future studies may constrain drawing time or use interactive tasks that incentivize prioritizing diagnostic scene components to provide better information on object prioritization within a scene.

In conclusion, our findings demonstrate that individual internal models shape gaze behavior when observers need to rely on expectations to guide information sampling. By linking personal expectations to exploration strategies, this work provides a new explanation of why individuals differ in how they look at and interpret visual environments. Incorporating idiosyncratic predictions into models of eye movements may advance computational accounts of attention and open new avenues for studying perception in real-world environments.

## Methods

### Participants

The study consisted of two experiments: free-viewing (Experiment 1) and gaze-contingent viewing (Experiment 2). For Experiment 1, we recruited 38 participants, of whom four were excluded due to insufficient eye-tracking calibration performance. This resulted in a sample size of 34 participants (gender distribution: 23 female, 11 male; mean age: 28.6 ± 5.1). For Experiment 2, we recruited 36 participants and excluded two, leaving also 34 participants (gender distribution: 20 female, 14 male; mean age: 26.1 ± 4.0). The sample size was selected to match previous studies that used drawings to capture inter-subject similarities in scene priors (Engeser & Kaiser, 2026). All participants reported normal (+/- 1.5 diopter) or corrected-to-normal vision, while only contact lenses were allowed for corrections. Participants were recruited through the participant recruitment system of the Justus Liebig University Giessen, provided written informed consent, and received monetary reimbursement of ten Euros per hour. Procedures were approved by the General Ethical Committee of the Justus Liebig University Giessen and adhered to the Declaration of Helsinki.

### Stimuli

The stimuli set consisted of 300 photographs of bathroom and kitchen scenes (150 each) selected from SUN database (Xiao et al., 2016) with LabelMe object mask annotations (Russell et al., 2008). Ten additional images (five per category) were chosen for a practice session, and another 150 (75 each) were selected as foils for the memory task in the gaze-contingency experiment. Presentation size was 19.9° x 15° visual angle at 68 cm viewing distance on a 32-inch Gigabyte AORUS FI32Q monitor (refresh rate: 60 Hz).

### Eye-tracking apparatus

Movements of both eyes were recorded at a frequency of 120 Hz using a monitor-mounted Tobii Pro Fusion eye tracker (TOBII, Stockholm, Sweden). The eye-tracker was calibrated using a nine-point calibration spanning 29.9° x 22.5° of visual angle and recalibrated after the practice block and every 100 images. Additional recalibrations could be performed, initiated by the experimenter.

### Procedure

The procedure of the two experiments was nearly identical, with all differences explicitly explained below. Participants first completed the drawing task (∼30 min), followed by the eye tracking task (∼90 min). In Experiment 2, participants performed an additional memory task (∼20 min). All experiments were coded and controlled in MATLAB R2022a (MathWorks, Natick, MA, USA) using the Psychtoolbox-3 (Kleiner et al., 2007) and Titta (Niehorster et al., 2020) toolboxes.

### Drawing Task

We used pictorial descriptions to assess individuals’ scene priors following a previously established pipeline (Engeser & Kaiser, 2025; Wang, Chen, et al., 2025; Wang, Duymaz, et al., 2025; Wang et al., 2024). Participants were asked to draw their most typical example of two scene categories (bathroom and kitchen). First, participants had 30 s to imagine the scene and four minutes to execute the drawing on an Apple iPad Pro (19.5 cm by 26.1 cm display) using an Apple Pencil, allowing them to produce black lines or erase them. A fixed template with the scene layout was provided at the beginning of each drawing to enforce a standard viewpoint. A video of the drawings was recorded to determine participants’ drawing order across object categories. After completing the drawings of typical scenes, participants additionally copied photographs of bathroom and kitchen scenes under the same four-minute time constraint. This served as a control condition capturing individual differences in drawing ability and style, as well as familiarization during the drawing process. This condition was later used to control for similarities in these unrelated factors during the IS-RSA.

### Eye-tracking task

In the eye-tracking task, participants viewed each of the 300 scene photographs once for three seconds in a random order that was fixed across participants (bathroom and kitchen images intermixed). Each trial started with a fixation cross presented at the center of the screen for at least 200 ms. Then participants could start stimulus presentation by pressing the space bar. To ensure central fixation at the start of each image presentation, the experiment only advanced if participants’ gaze (average of both eyes) fell within a central square of 1.3° visual angle at the moment of the button press. This also served as a quality check of eye-tracker calibration: When the experiment did not advance to the next image while the participant reported central fixation, a recalibration was initiated by the experimenter. In a brief practice session (ten and six images for Experiment 1 and Experiment 2, respectively), participants could familiarize themselves with the setup and procedures. A mandatory break was added after every 100 images.

In Experiment 1, participants were asked to freely view the images without further task constraints, and images were presented in full resolution and color. By contrast, in Experiment 2, only a small, circular window of 2.5° visual angle radius around the current gaze position (average of both eyes) was shown at full resolution, while the rest of the image was heavily degraded (2-D Gaussian smoothing with a 20 standard deviation kernel and desaturation by eliminating hue and saturation information while retaining luminance). To ensure smooth movement of the gaze-contingent window, we added a Kalman filter (0.01 process noise and 0.1 measurement noise) and an update threshold of 0.5° visual angle (position of the gaze-contingent window was only updated when the distance between the current gaze-position and the previous window position exceeded the threshold). Moreover, participants were instructed to explore the images carefully, so they would remember them well in a subsequent memory task.

### Memory task

Experiment 2 also included a memory task after the eye-tracking session, consisting of 150 trials. In each trial, a pair of images was presented (29.9° x 22.5° of visual angle image-size and 16.9° eccentricity). One of the images was previously presented in the eye-tracking experiment, while the other one was novel. For each of 150 images (75 bathroom and 75 kitchen scenes) randomly selected from the eye-tracking experiment, a roughly similar-looking novel foil from the same scene category was hand-selected from the SUN Database. Participants indicated which image they had seen during the eye-tracking task. The position of the correct image was randomized. Participants received immediate feedback after each trial. Image selection and trial randomization were performed once and kept fixed across participants.

### Analysis

For estimating the relationship between individual differences in scene priors and gaze exploration patterns, we employed an inter-subject representational similarity analysis (IS-RSA) framework (Finn et al., 2020). For each variable of interest, such as drawings and gaze measures, we constructed an inter-subject representational dissimilarity matrix (IS-RDM) of size n x n (where n is the sample size of the respective experiment) representing a common similarity space that allows for comparing the inter-subject variance across different variables.

All analysis steps were performed separately for the two scene categories, and only the final correlation values were averaged before statistical testing.

### Inter-subject similarities in drawings

Firstly, we assessed individual differences in participants’ scene priors using the relatively unconstrained, descriptive measure of drawings. To objectively estimate the similarity between these drawings, we followed the analysis pipeline employed in Engeser & Kaiser (2025). In brief, we used correlations of the feature activations of a deep neural network (DNN, penultimate layer “fc7” of VGG-16 (Simonyan & Zisserman, 2015) pre-trained on ImageNet (Deng et al., 2009)) to approximate high-level, perceptual similarity in humans. Instead of directly presenting the drawings to the DNN, we used a diffusion-based AI method (commercial Draw3D software; https://draw3d.online) to generate photorealistic images from the drawings, which align better with the training material of the DNN. Three images were generated for each drawing (to account for variance across image generations) and converted to greyscale. Image generation was completed before the inspection of subsequent analysis results. Next, an IS-RDM was built using the dissimilarities (1 - Spearman R) for each pair of participants (averaging the correlation coefficients across the triplets of generated images), representing inter-subject dissimilarities in individuals’ internal scene models.

### Inter-subject similarities in scene exploration responses

To quantify individual differences in gaze, we measured how participants differed in (i) gaze distance, (ii) fixation counts, (iii) object dwell times, and (iv) fixation order.

The calculation of gaze distance used raw gaze positions (averaged between eyes) while the remaining analysis approaches were fixation-based measures. For these analyses, fixations were classified using the MATLAB implementation of the I2MC algorithm (Hessels et al., 2017) with default settings for 120 Hz sampling frequency. Moreover, fixations shorter or with an onset earlier than 100 ms were excluded from the analysis.

First, we employed a straightforward distance measure to estimate how people differ in their exploration trajectories for natural scenes. Thereby, the median Euclidean distance of gaze positions for each pair of subjects and each image was calculated and then averaged across all images of the same category, resulting in an IS-RDM of gaze distances. From a simple distance-based measure, it is hard to disentangle whether differences between individuals emerge from looking at different objects or whether they simply look at them in a different order. Therefore, we used three additional measures to dissociate these aspects.

Second, to account for the different dynamics of scene exploration, we counted the number of fixations each subject executed for each image. An IS-RDM was then created by calculating the pairwise dissimilarities (1 - Spearman correlation coefficients) in fixation counts across all images.

Third, we tested how people differ in how long they look at which kind of objects. Therefore, we measured the proportion of time the fixations fell within a binary object mask generated from LabeME scene annotations (Russell et al., 2008). To account for minor imprecisions in measuring the gaze positions, we enlarged all masks by 0.5° visual angle. All individual annotated object labels were grouped into coarse object categories (i.e., “shower head”, “shower screen”, and “shower curtain” were grouped into the “shower” category). We restricted our analysis to object categories that appeared in at least 10 images per scene category and removed background labels (i.e., “floor”, “ceiling”, and “wall”), leaving 35 object categories for bathrooms and 66 categories for kitchens. For each image, we accumulated fixation durations that fell on objects belonging to the same category and divided them by the sum of all fixations on the respective image. In case of overlapping object masks (i.e., a “pot” label may overlap with the label of “stove”), fixations in the overlapping area were attributed to both object categories. Then, the proportional dwell times were summed for each object category across all images of the same scene category, and an IS-RDM was created by calculating the pairwise dissimilarities (1 - Spearman correlation coefficients) in the resulting dwell times across object categories.

Finally, we analyzed the order in which objects were looked at. Therefore, we estimated a priority score for each object category representing the time it takes for an object of that category to be looked at. For all images that featured a respective object category, we extracted the first time an object of that category was fixated and averaged that value across images. When an object category was not fixated despite being present in an image, the full three seconds were assigned for that object category for that specific image. Then, an IS-RDM was created by calculating the pairwise dissimilarities (1 - Spearman correlation coefficients) in priority scores across all object categories.

### Split-half reliability for individual differences in gaze behavior

To assess the reliability of individual differences in the gaze pattern, we conducted a split-half reliability analysis. Therefore, the four measures of gaze-similarity were calculated separately for odd and even images, resulting in two IS-RDMs for each measure based on independent image sets. A positive correlation between these two IS-RDMs provides evidence for reliable individual differences captured with the respective measures. The split-half correlation coefficients were computed separately for the two scene categories and then averaged.

### Comparison of inter-subject similarities in drawings and gaze patterns

To test our hypothesis that participants with more similar scene priors explore a scene in more similar ways, we correlated the IS-RDMs for each gaze measure with the IS-RDMs constructed from the typical drawings while partialling out the IS-RDMs constructed from the copy control drawings. Specifically, we used Pearson correlations of the off-diagonal values in the lower triangle of the IS-RDMs separately for each of the two scene categories and averaged the resulting correlation coefficient.

### Comparison of drawing order and fixation order

We tested whether the order in which the objects are placed in the drawing reveals information about their priority in people’s scene priors and whether this can predict individual differences in the gaze trajectory during scene exploration. The drawing order of the object categories identified above was extracted from the videos of the typical scene drawings. We focused on the first stroke of the first object that belonged to a respective object category, irrespective of when the drawing of that object was completed.

We then assessed whether a subject’s drawing order better predicts that subject’s own fixation order compared to other subjects’ fixation orders. For each subject, we selected the fixation priority scores of the object categories that a participant drew and correlated (Spearman) them with their drawing order. Then the fixation priority scores of the same object categories for all other subjects were correlated with that participant’s drawing order, and the mean of the resulting correlation coefficients was subtracted from the coefficients from the correlation with their own fixation order. A difference (Δr) greater than zero indicated that a participant’s drawing order better predicts their own fixation order than that of other participants. Lastly, the differences were averaged across all subjects.

### Memory task performance

The memory task in the gaze-contingency experiment tested whether participants could distinguish previously presented images from novel foils. Accuracy for each participant was computed as the proportion of correctly remembered images. The mean accuracy was 58% (SD=8%, p<0.001), indicating that the memory task was challenging but that participants could remember the viewed scenes above chance (50%), despite receiving limited, gaze-contingent scene information.

We then tested whether individual differences in the memory task performance are related to individual differences in internal models or viewing behaviors. A memory task IS-RDM was built by correlating (Phi) the accuracy scores across all memory task trials of the same scene category between each pair of observers. Correlations of that memory task IS-RDM with IS-RDMs of the four gaze measures were not significant (gaze distance: r=-0.02; fixation count: r=0.03; object dwells: r=0.04; fixation order: r=0.06; all p>0.31). Also, the inter-subject similarities in memory performance could not be predicted by inter-subject similarities in internal models (r=0.06; all p=0.12).

### Inter-subject similarities across experiments

To test whether the gaze-contingency manipulation in Experiment 2 substantially altered gaze behavior, we compared inter-subject similarities within and across experiments. IS-RDMs for all four gaze measures were constructed using participants from both experiments combined. We then compared the median pairwise (dis)similarities between participant pairs within the same experiment and between pairs drawn from different experiments.

For the gaze distance measure, we subtracted the median distance between participants from the same experiment from the median distance between participants from different experiments, testing whether gaze positions were more similar within the same experimental condition. For the remaining measures (number of fixations per image, object dwell time, and fixation order), which were based on inter-subject correlations, we computed the difference between within-experiment and across-experiment correlations to test whether gaze similarity was higher among participants who performed the same experiment.

### Statistical analysis

To assess statistical significance in the partial correlation of gaze and drawing IS-RDMs or the split-half correlations, we used a non-parametric subject-wise permutation test (Chen et al., 2016). The partial correlation was repeated 10,000 times, while at each iteration the rows and columns of the gaze IS-RDM were randomly shuffled. At each iteration, the correlations were conducted separately for the two scene categories and then averaged. Employing a one-sided test against zero, the p-value is obtained by determining the proportion of permutations that result in equal or higher correlation coefficients than the correlation of interest. False discovery rate (FDR) correction was applied across gaze similarity measures using the Benjamini and Hochberg procedure (Benjamini & Hochberg, 1995).

In the memory task, accuracy values across participants were compared against chance level accuracy (50%) using a one-sided t-test.

In the comparison of drawing with fixation order, significance was determined by a one-sided t-test of the differences in own-vs-other correlations against zero.

For the comparison across experiments, a permutation test was used, in which the within-and across-experimental inter-subject (dis)similarities were compared 10,000 times while randomly shuffling the assignment of participants between the two experiments in each iteration.

## Data and Code availability

All data and code are freely available from GitHub (https://github.com/DKaiserLab/pep_wp4_eye_tracking). The photos of participants’ private rooms and raw anatomical scans are not shared to protect participants’ privacy. The experiment and analysis are implemented in MATLAB (R2022a), including Psychtoolbox-3 and Tittta toolboxes. Detailed instructions for installation and execution of the code can be found in the corresponding README file.

## Acknowledgements

This work was supported by the Deutsche Forschungsgemeinschaft (DFG), grants KA4683/5-1 (project no. 518483074), KA4683/7-1 (project no. 548389777), and under Germany’s Excellence Strategy (EXC 3066/1, “The Adaptive Mind”, project no. 533717223). It was further supported by a European Research Council (ERC) Starting Grant (PEP, ERC-2022-STG 101076057). Views and opinions expressed are those of the authors only and do not necessarily reflect those of the funders. Neither the funders nor the granting authority can be held responsible for them.

We thank Bati Yilmaz for providing feedback on the manuscript. The authors declare that there are no competing interests.

## Supplement

**Figure S1.**
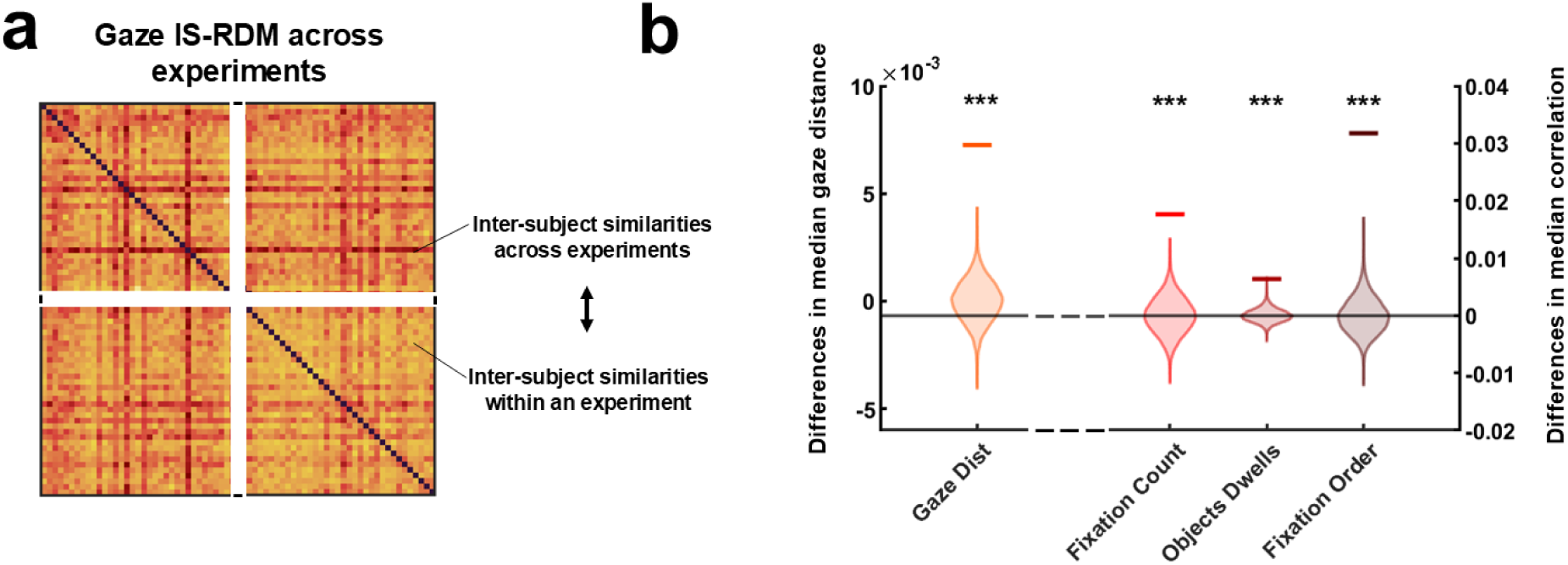
Comparison across experiments. **(a)** IS-RDMs for all four gaze measures were constructed with the subjects of both experiments combined. We compared the median (dis)similarities between participants of the same experiment and between experiments. **(b)** The median gaze distance was smaller between individuals within the same experiment than across experiments. Similarly, the inter-subject correlation in the number of fixations, the object dwell times, and fixation order was higher when participants performed the same experiment. Horizontal bars indicate observed correlation, and violin plots depict the empirical null distribution from the permutation test. *** p<0.001.

## References

Andrews, T. J., & Coppola, D. M. (1999). Idiosyncratic characteristics of saccadic eye movements when viewing different visual environments. Vision Research, 39(17), 2947–2953. 10.1016/S0042-6989(99)00019-X

Bakst, L., & McGuire, J. T. (2021). Eye movements reflect adaptive predictions and predictive precision. Journal of Experimental Psychology: General, 150(5), 915–929. 10.1037/xge0000977

Bargary, G., Bosten, J. M., Goodbourn, P. T., Lawrance-Owen, A. J., Hogg, R. E., & Mollon, J. D. (2017). Individual differences in human eye movements: An oculomotor signature? *Vision Research*, Individual Differences as a Window into the Structure and Function of the Visual System, 141, 157–169. 10.1016/j.visres.2017.03.001

Bartnik, C. G., Sartzetaki, C., Sanchez, A. P., Molenkamp, E., Bommer, S., Vukšić, N., & Groen, I. I. A. (2025). Representation of locomotive action affordances in human behavior, brains, and deep neural networks. Proceedings of the National Academy of Sciences, 122(24), e2414005122. 10.1073/pnas.2414005122

Benjamini, Y., & Hochberg, Y. (1995). Controlling the False Discovery Rate: A Practical and Powerful Approach to Multiple Testing. Journal of the Royal Statistical Society: Series B (Methodological), 57(1), 289–300. 10.1111/j.2517-6161.1995.tb02031.x

Borovska, P., & de Haas, B. (2024). Individual gaze shapes diverging neural representations. Proceedings of the National Academy of Sciences, 121(36), e2405602121. 10.1073/pnas.2405602121

Broda, M. D., & de Haas, B. (2024). Individual differences in human gaze behavior generalize from faces to objects. Proceedings of the National Academy of Sciences, 121(12), e2322149121. 10.1073/pnas.2322149121

Brunyé, T. T., & Gardony, A. L. (2017). Eye tracking measures of uncertainty during perceptual decision making. International Journal of Psychophysiology, 120, 60–68. 10.1016/j.ijpsycho.2017.07.008

Charest, I., Kievit, R. A., Schmitz, T. W., Deca, D., & Kriegeskorte, N. (2014). Unique semantic space in the brain of each beholder predicts perceived similarity. Proceedings of the National Academy of Sciences, 111(40), 14565–14570. 10.1073/pnas.1402594111

Chen, G., Shin, Y.-W., Taylor, P. A., Glen, D. R., Reynolds, R. C., Israel, R. B., & Cox, R. W. (2016). Untangling the relatedness among correlations, part I: Nonparametric approaches to inter-subject correlation analysis at the group level. NeuroImage, 142, 248–259. 10.1016/j.neuroimage.2016.05.023

Coco, M. I., & Keller, F. (2012). Scan Patterns Predict Sentence Production in the Cross-Modal Processing of Visual Scenes. Cognitive Science, 36(7), 1204–1223. 10.1111/j.1551-6709.2012.01246.x

Constantino, J. N., Kennon-McGill, S., Weichselbaum, C., Marrus, N., Haider, A., Glowinski, A. L., Gillespie, S., Klaiman, C., Klin, A., & Jones, W. (2017). Infant viewing of social scenes is under genetic control and is atypical in autism. Nature, 547(7663), 340–344. 10.1038/nature22999

de Haas, B., Iakovidis, A. L., Schwarzkopf, D. S., & Gegenfurtner, K. R. (2019). Individual differences in visual salience vary along semantic dimensions. Proceedings of the National Academy of Sciences, 116(24), 11687–11692. 10.1073/pnas.1820553116

de Lange, F. P., Heilbron, M., & Kok, P. (2018). How Do Expectations Shape Perception? Trends in Cognitive Sciences, 22(9), 764–779. 10.1016/j.tics.2018.06.002

Deng, J., Dong, W., Socher, R., Li, L.-J., Kai Li, & Li Fei-Fei. (2009). ImageNet: A large-scale hierarchical image database. IEEE Conference on Computer Vision and Pattern Recognition, 248–255. 10.1109/CVPR.2009.5206848

Ekinci, M. A., & Kaiser, D. (2026). Shared gaze reflects shared aesthetic experiences (p. 2026.01.30.702749). bioRxiv. 10.64898/2026.01.30.702749

Engeser, M., Ajith, S., Duymaz, I., Wang, G., Foxwell, M. J., Cichy, R. M., Pitcher, D., & Kaiser, D. (2025). Characterizing internal models of the visual environment. Proceedings of the Royal Society B: Biological Sciences, 292(2053), 20250602. 10.1098/rspb.2025.0602

Engeser, M., & Kaiser, D. (2025). Inter-individual similarities in internal models of the world shape similarities in the perception and neural processing of scenes. OSF. 10.17605/OSF.IO/ZJTWX

Finn, E. S., Glerean, E., Khojandi, A. Y., Nielson, D., Molfese, P. J., Handwerker, D. A., & Bandettini, P. A. (2020). Idiosynchrony: From shared responses to individual differences during naturalistic neuroimaging. NeuroImage, 215, 116828. 10.1016/j.neuroimage.2020.116828

Friston, K., Adams, R., Perrinet, L., & Breakspear, M. (2012). Perceptions as Hypotheses: Saccades as Experiments. Frontiers in Psychology, 3. 10.3389/fpsyg.2012.00151

Friston, K., Daunizeau, J., Kilner, J., & Kiebel, S. J. (2010). Action and behavior: A free-energy formulation. Biological Cybernetics, 102(3), 227–260. 10.1007/s00422-010-0364-z

Goettker, A., Pidaparthy, H., Braun, D. I., Elder, J. H., & Gegenfurtner, K. R. (2021). Ice hockey spectators use contextual cues to guide predictive eye movements. Current Biology, 31(16), R991–R992. 10.1016/j.cub.2021.06.087

Greene, M. R., Baldassano, C., Esteva, A., Beck, D. M., & Fei-Fei, L. (2016). Visual scenes are categorized by function. Journal of Experimental Psychology: General, 145(1), 82–94. 10.1037/xge0000129

Han, C., & Bonner, M. F. (2026). High-dimensional structure underlying individual differences in naturalistic visual experience. Current Biology, 36(3), 723–733.e6. 10.1016/j.cub.2025.12.039

Hartley, C. A. (2022). How do natural environments shape adaptive cognition across the lifespan? Trends in Cognitive Sciences, 26(12), 1029–1030. 10.1016/j.tics.2022.10.002

Hayes, T. R., & Henderson, J. M. (2017). Scan patterns during real-world scene viewing predict individual differences in cognitive capacity. Journal of Vision, 17(5), 23. 10.1167/17.5.23

Hayhoe, M. M., McKinney, T., Chajka, K., & Pelz, J. B. (2012). Predictive eye movements in natural vision. Experimental Brain Research, 217(1), 125–136. 10.1007/s00221-011-2979-2

Henderson, J. M. (2017). Gaze Control as Prediction. Trends in Cognitive Sciences, 21(1), 15–23. 10.1016/j.tics.2016.11.003

Henderson, J. M., & Luke, S. G. (2014). Stable individual differences in saccadic eye movements during reading, pseudoreading, scene viewing, and scene search. Journal of Experimental Psychology: Human Perception and Performance, 40(4), 1390–1400. 10.1037/a0036330

Hessels, R. S., Niehorster, D. C., Kemner, C., & Hooge, I. T. C. (2017). Noise-robust fixation detection in eye movement data: Identification by two-means clustering (I2MC). Behavior Research Methods, 49(5), 1802–1823. 10.3758/s13428-016-0822-1

Itti, L., & Koch, C. (2001). Computational modelling of visual attention. Nature Reviews Neuroscience, 2(3), 194–203. 10.1038/35058500

Itti, L., Koch, C., & Niebur, E. (1998). A model of saliency-based visual attention for rapid scene analysis. IEEE Transactions on Pattern Analysis and Machine Intelligence, 20(11), 1254–1259. 10.1109/34.730558

Kennedy, D. P., D’Onofrio, B. M., Quinn, P. D., Bölte, S., Lichtenstein, P., & Falck-Ytter, T. (2017). Genetic Influence on Eye Movements to Complex Scenes at Short Timescales. Current Biology, 27(22), 3554–3560.e3. 10.1016/j.cub.2017.10.007

Kleiner, M., Brainard, D., & Pelli, D. (2007). What’s new in Psychtoolbox*-*3*?*

Kollenda, D., Reher, A.-S., & de Haas, B. (2025). Individual gaze predicts individual scene descriptions. Scientific Reports, 15(1), 9443. 10.1038/s41598-025-94056-4

Lee, J., & Geng, J. J. (2017). Idiosyncratic Patterns of Representational Similarity in Prefrontal Cortex Predict Attentional Performance. The Journal of Neuroscience: The Official Journal of the Society for Neuroscience, 37(5), 1257–1268. 10.1523/JNEUROSCI.1407-16.2016

Linka, M., & de Haas, B. (2026). Detection, Inspection, Return: An Object-Based Classification and Metric of Fixations in Complex Scenes. Open Mind, 10, 47–65. 10.1162/OPMI.a.319

Linka, M., Karimpur, H., & de Haas, B. (2025). Protracted development of gaze behaviour. Nature Human Behaviour, 9(9), 1887–1897. 10.1038/s41562-025-02191-9

Malcolm, G. L., & Henderson, J. M. (2010). Combining top-down processes to guide eye movements during real-world scene search. Journal of Vision, 10(2), 4. 10.1167/10.2.4

Mollon, J. D., Bosten, J. M., Peterzell, D. H., & Webster, M. A. (2017). Individual differences in visual science: What can be learned and what is good experimental practice? *Vision Research*, Individual Differences as a Window into the Structure and Function of the Visual System, 141, 4–15. 10.1016/j.visres.2017.11.001

Murlidaran, S., & Eckstein, M. P. (2025). Eye movements during free viewing to maximize scene understanding. Nature Communications, 17(1), 940. 10.1038/s41467-025-67673-w

Najemnik, J., & Geisler, W. S. (2005). Optimal eye movement strategies in visual search. Nature, 434(7031), 387–391. 10.1038/nature03390

Najemnik, J., & Geisler, W. S. (2008). Eye movement statistics in humans are consistent with an optimal search strategy. Journal of Vision, 8(3), 4. 10.1167/8.3.4

Niehorster, D. C., Andersson, R., & Nyström, M. (2020). Titta: A toolbox for creating PsychToolbox and Psychopy experiments with Tobii eye trackers. Behavior Research Methods, 52(5), 1970–1979. 10.3758/s13428-020-01358-8

Palmer, S. E., Schloss, K. B., & Sammartino, J. (2013). Visual Aesthetics and Human Preference. Annual Review of Psychology, 64(Volume 64, 2013), 77–107. 10.1146/annurev-psych-120710-100504

Peelen, M. V., Berlot, E., & de Lange, F. P. (2024). Predictive processing of scenes and objects. Nature Reviews Psychology, 3(1), Article 1. 10.1038/s44159-023-00254-0

Richler, J. J., Tomarken, A. J., Sunday, M. A., Vickery, T. J., Ryan, K. F., Floyd, R. J., Sheinberg, D., Wong, A. C.-N., & Gauthier, I. (2019). Individual differences in object recognition. Psychological Review, 126(2), 226–251. (2019-09750-002). 10.1037/rev0000129

Rothkegel, L. O. M., Schütt, H. H., Trukenbrod, H. A., Wichmann, F. A., & Engbert, R. (2018). *Searchers adjust their eye movement dynamics to the target characteristics in natural scenes* (arXiv:1802.04069). arXiv. 10.48550/arXiv.1802.04069

Russell, B. C., Torralba, A., Murphy, K. P., & Freeman, W. T. (2008). LabelMe: A Database and Web-Based Tool for Image Annotation. International Journal of Computer Vision, 77(1), 157–173. 10.1007/s11263-007-0090-8

Schwetlick, L., Backhaus, D., & Engbert, R. (2023). A dynamical scan-path model for task-dependence during scene viewing. Psychological Review, 130(3), 807–840. 10.1037/rev0000379

Sclar, M., Bujia, G., Vita, S., Solovey, G., & Kamienkowski, J. E. (2020). *Modeling human visual search: A combined Bayesian searcher and saliency map approach for eye movement guidance in natural scenes* (arXiv:2009.08373). arXiv. 10.48550/arXiv.2009.08373

Simonyan, K., & Zisserman, A. (2015). *Very Deep Convolutional Networks for Large-Scale Image Recognition* (arXiv:1409.1556). arXiv. 10.48550/arXiv.1409.1556

Singer, J. J. D., Cichy, R. M., & Hebart, M. N. (2023). The Spatiotemporal Neural Dynamics of Object Recognition for Natural Images and Line Drawings. Journal of Neuroscience, 43(3), 484–500. 10.1523/JNEUROSCI.1546-22.2022

Spotorno, S., Malcolm, G. L., & Tatler, B. W. (2014). How context information and target information guide the eyes from the first epoch of search in real-world scenes. Journal of Vision, 14(2), 7–7.

Vanmarcke, S., & Wagemans, J. (2016). Individual differences in spatial frequency processing in scene perception: The influence of autism-related traits. Visual Cognition, 24(2), 115–131. 10.1080/13506285.2016.1199625

Vessel, E. A., Maurer, N., Denker, A. H., & Starr, G. G. (2018). Stronger shared taste for natural aesthetic domains than for artifacts of human culture. Cognition, 179, 121–131. 10.1016/j.cognition.2018.06.009

Võ, M. L.-H., Boettcher, S. E., & Draschkow, D. (2019). Reading scenes: How scene grammar guides attention and aids perception in real-world environments. *Current Opinion in Psychology*, Attention & Perception, 29, 205–210. 10.1016/j.copsyc.2019.03.009

Wang, G., Chen, L., Cichy, R. M., & Kaiser, D. (2025). Enhanced and idiosyncratic neural representations of personally typical scenes. Proceedings of the Royal Society B: Biological Sciences, 292(2043), 20250272. 10.1098/rspb.2025.0272

Wang, G., Duymaz, I., Foxwell, M., Engeser, M., Pitcher, D., Cichy, R. M., & Kaiser, D. (2025). How do visual and conceptual factors predict the composition of typical scene drawings? (p. 2025.09.15.676247). bioRxiv. 10.1101/2025.09.15.676247

Wang, G., Foxwell, M. J., Cichy, R. M., Pitcher, D., & Kaiser, D. (2024). Individual differences in internal models explain idiosyncrasies in scene perception. Cognition, 245, 105723. 10.1016/j.cognition.2024.105723

Xiao, J., Ehinger, K. A., Hays, J., Torralba, A., & Oliva, A. (2016). SUN Database: Exploring a Large Collection of Scene Categories. International Journal of Computer Vision, 119(1), 3–22. 10.1007/s11263-014-0748-y

Yun, K., Peng, Y., Samaras, D., Zelinsky, G. J., & Berg, T. L. (2013). Exploring the role of gaze behavior and object detection in scene understanding. Frontiers in Psychology, 4. 10.3389/fpsyg.2013.00917

